# Predict plant-derived xenomiRs from plant miRNA sequences using random forest and one-dimensional convolutional neural network models

**DOI:** 10.1101/345249

**Authors:** Qi Zhao, Qian Mao, Zheng Zhao, Tongyi Dou, Zhiguo Wang, Xiaoyu Cui, Yuanning Liu, Xiaoya Fan

## Abstract

**Background:** An increasing number of studies reported that exogenous miRNAs (xenomiRs) can be detected in animal bodies, however, some others reported negative results. Some attributed this divergence to the selective absorption of plant-derived xenomiRs by animals.

**Results:** Here, we analyzed 166 plant-derived xenomiRs reported in our previous study and 942 non-xenomiRs extracted from miRNA expression profiles of four species of commonly consumed plants. Employing statistics analysis and cluster analysis, our study revealed the potential sequence specificity of plant-derived xenomiRs. Furthermore, a random forest model and a one-dimensional convolutional neural network model were trained using miRNA sequence features and raw miRNA sequences respectively and then employed to predict unlabeled plant miRNAs in miRBase. A total of 241 possible plant-derived xenomiRs were predicted by both models. Finally, the potential functions of these possible plant-derived xenomiRs along with our previously reported ones in human body were analyzed.

**Conclusions:** Our study, for the first time, presents the systematic plant-derived xenomiR sequences analysis and provides evidence for selective absorption of plant miRNA by human body, which could facilitate the future investigation about the mechanisms underlying the transference of plant-derived xenomiR.

## Background

miRNAs and their gene expression regulation function in eukaryotes is one of the most important discoveries in recent years [1]. It has been well elucidated that endogenous miRNAs could degrade or silence mRNAs to mediate gene expression by binding RNA-induced silencing complex (RICS) in a sequence-specific manner [2]. Meanwhile, although still controversial, new hypotheses about extracellular miRNA have been continually proposed, e.g., exosomal miRNA [3, 4], circulating miRNA [5, 6] and exogenous miRNA (xenomiR) [7, 8].

Due to the possibility of cross-kingdom regulation, plant-derived xenomiR hypothesis has received great attention since first proposed in 2012 [7]. Plant-derived xenomiRs were defined as the miRNAs derived from plants which are capable of transferring into human or animal bodies. Subsequently, plant-derived xenomiRs have been detected in different tissues or body fluids of several species of animals, including human [7], mice [7], pig [9], panda [10] and silkworm[11]. And their relevance to many diseases, such as cardiovascular diseases [7], tumor [12, 13], chronic-inflammation [14], influenza [15], benign prostatic hyperplasia [16] and pulmonary fibrosis [17], were also proposed. However, many mechanisms of plant-derived xenomiRs in keeping stable in gastrointestinal (GI) track, transferring across GI track, entering cells or being secreted by cells are still unknown.

Emerging evidence suggests that the species of plant miRNAs detected in animals are limited, although the total species of miRNAs of a single plant is often more than several hundred, for example 713 species of miRNAs have been identified in *oryza sativa* (osa) so far [18]. Only 25 species of plant miRNAs were detected by Zhang et al. [7], although their samples pooled 80 human serum (8 samples, each sample pooled from 10 humans). In another study of Zhang et al. [19], where plant miRNAs in human plasma were examined by qRT-PCR after donors drunk fruit juice, 10 species of plant miRNAs were detected, whereas 16 species of plant miRNAs could be detected in the fruit juice. Similarly, limited species of maize miRNAs were detected using qRT-PCR in the serum and tissues of pigs feed with fresh maize for 7 days [20]. With TA-cloning and Sanger sequencing, only a part of species of mulberry-derived miRNAs were detected in hemolymphs of silkworms which were fed with mulberry leaves [11].

In fact, besides natural plant miRNAs, many species of synthetic plant miRNAs or mimic plant miRNAs that are identical or similar to natural plant miRNAs, were also reported to be able to transfer into and keep stable in animal bodies. Chin et al. [12] suggested that both natural plant miR159 and synthetic oligos, with the same sequence as miR159, were capable of transferring into human breast cancer cells. Similarly, the synthetic miR166b were detected in silkworm hemolymph [11]. A recent study reported that 3 species of mimic plant miRNAs (mmu-miR34a, mmu-miR143 and mmu-miR145) can also transfer into mouse body by oral administration [13]. Many reports suggested that MIR2911 could be significantly taken in by human and animals, which was attributed to its unique sequence [15, 21, 22], and the disruption of the MIR2911 sequence by two nucleotides abolished its absorption [23]. Yang et al. [22] suggested that not all miRNAs, but miRNAs with certain features could keep stable in GI tract of animals, and randomly synthesized miRNA-like sequences would be degraded quickly after injected in animals.

The discoveries described above imply the selective absorption of plant miRNAs by animals, i.e. only plant miRNAs with specific sequence could be absorbed by specific species of animals, which also provides an explanation for studies that reported the un-detectability [24, 25] of several plant miRNAs in animals. In this paper, we first systematically studied the sequence differences between the plant miRNAs which can transfer (xenomiR group) and cannot transfer (non-xenomiR group) into human bodies using statistics methods. Significant difference was found in 28 sequence features between the two groups, which suggested the potential patterns underlying the plant-derived xenomiR sequences and a possible link between these patterns with selective absorption of xenomiRs. Subsequently, a random forest (RF) model and a onedimensional convolutional neural network (1D-CNN) model were trained to distinguish the two groups. Both models successfully distinguished between xenomiRs and non-xenomiRs with high accuracy. They were then used to predict potential plant-derived xenomiRs on unlabeled plant miRNAs, and a total of 241 plant miRNAs were identified as xenomiRs by both models. Finally, we analyzed the functions of the 241 predicted along with 166 previously reported xenomiRs in human body. Taken together, we report the first systematic plant-derived xenomiR sequences analysis, and the results provide evidence for selective absorption of plant miRNAs by human body. In addition, we propose the first list of high-probability xenomiRs, which further enables more robust decisions regarding plant miRNAs candidates for experimental validation and facilitates future investigation about the mechanisms of xenomiRs transferring into animal bodies.

## Results

### Datasets and feature extraction

For sequence comparison, we collected 166 xenomiR sequences (positive samples) and 942 non-xenomiR sequences (negative samples). All 166 xenomiRs (Additional file 2: Table S1) were collected from our previous study [26], which were obtained from 388 healthy human samples analyzed by a rigorous bioinformatics pipeline. These miRNAs covered almost all the reported plant-derived xenomiRs so far. Regarding non-xenomiRs (negative samples), no off the shelf dataset is available at present. To obtain the non-xenomiRs as accurately as possible, we carefully selected the miRNAs that have never been detected in human from *osa, zea maize* (*zma*)*, glycine max* (*gma*) and *arabidopsis thaliana* (*ath*) (see Methods), which are either staple food or the plant closely related to common vegetables (see Discussion). In total, 942 miRNAs (Additional file 3: Table S2) were labeled as non-xenomiRs. For both positive and negative samples, we extracted the length, nucleotide positions, 1^~^3 nt motif frequency in full miRNA sequences and 1^~^2 nt motif frequency in miRNA seed regions (Additional file 4: Table S3). All these features are widely used in miRNA associated researches.

### Statistical analysis of the differences between xenomiRs and non-xenomiRs

Considering the similarity of the members in the same miRNA family, we first studied the miRNA families to which the xenomiRs belong according to the miRNA families classified by miRBase [18]. In total, 49 miRNA families were mapped by xenomiRs (Additional file 5: Table S4), among which 8 (Fig 5-xenomiR family) covered nearly half of the 166 xenomiRs. In mir168 family, up to 41.2% miRNAs (7 out of 17 miRNA sequences) were mapped by xenomiRs. These results suggested that xenomiRs are likely to enrich in specific miRNA families, rather than randomly distributed among all miRNA families.

Further, we explored the differences between xenomiRs (Additional file 2: Table S1) and non-xenomiRs (Additional file 3: Table S2) in terms of nucleotide position. The percentages of the 4 kinds of nucleotides (adenine (A), cytosine (C), guanine (G) and uracil (U)) in each position were obtained for each group (Fig 2-position comparison), respectively. It can be found that the percentages of nucleotides are different in xenomiRs and non-xenomiRs, especially the percentages of pyrimidines (U at the 1^st^, 7^th^, 9^th^, 13^th^, 15^th~^17^th^ position, and C at the 3^rd^, 4^th^, 5^th^,6^th^, 9^th^, 13^th^, 15^th~^22^nd^ position), suggesting the difference in position features between the two groups. Flypothesis tests were also performed on all the other features listed in Additional file 4: Table S3 for further comparison. In total, 28 out of 105 features were significantly different between the two groups (p < 0.05, false discovery rate (FDR) corrected), as listed in Table 1-feature-comparison, including length, C content, U content, 4 kinds of 2 mer motif, 17 kinds of 3 mer motif and 4 kinds of 2 mer motif in seed region. It can be found that, comparing with non-xenomiRs, the contents of most motifs (1^~^3 mer) with C nucleotides are higher in xenomiRs, yet the contents of most motifs with U nucleotides are lower. Besides, the sequence length of xenomiRs is also significantly shorter than that of non-xenomiRs. Taken together, our results suggested that xenomiRs and non-xenomiRs are separable in sequence feature space.

**Fig 1.**
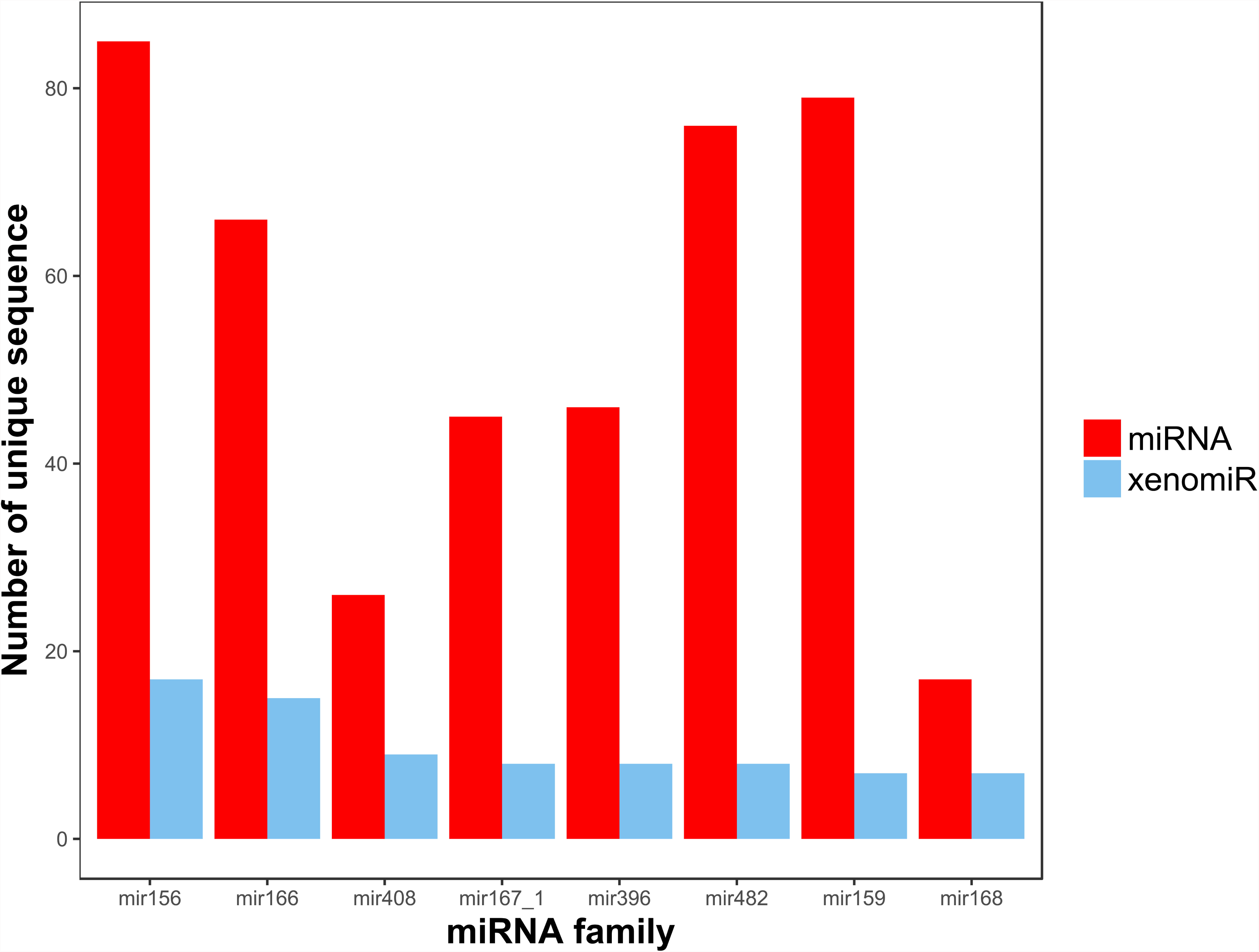
The top 8 plant miRNA families that contain the most xenomiRs. About one half of xenomiRs belong to these 8 miRNA families. In mir168 family, up to 41.2% miRNAs (7 of 17 miRNA sequences) were mapped.

**Fig 2.**
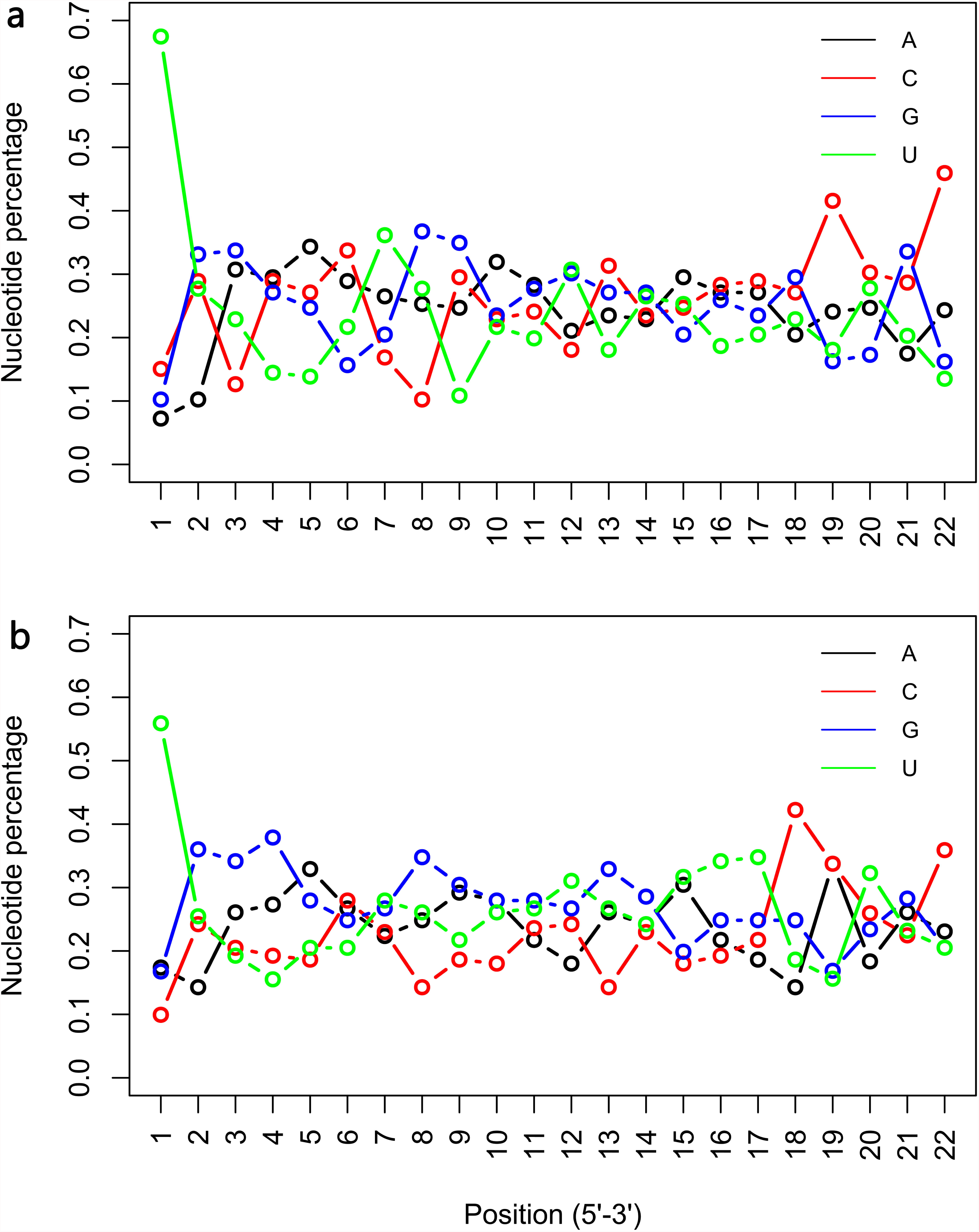
Nucleotide position comparison between xenomiRs and non-xenomiRs. Percentage of the four kinds of nucleotide at each position of a) 166 xenomiRs and b) 942 non-xenomiRs.

**Table 1.** Sequence feature comparison between xenomiRs and non-xenomiRs

| Feature | p value (t-test) | FDR | Mean (positive) | Mean (negative) |
| --- | --- | --- | --- | --- |
| Length | 2.47E-08 | 5.20E-07 | 21.07229 | 21.44586 |
| **C** | **2.76E-06** | **2.89E-05** | **0.2547** | **0.206519** |
| U | 1.38E-04 | 7.23E-04 | 0.242166 | 0.276587 |
| AU | 2.01E-05 | 1.41E-04 | 0.051751 | 0.068726 |
| **CA** | **9.01E-09** | **2.37E-07** | **0.085152** | **0.060352** |
| GU | 8.07E-09 | 2.37E-07 | 0.034289 | 0.057152 |
| UA | 4.83E-14 | 2.54E-12 | 0.021754 | 0.045344 |
| **AGC** | **5.19E-04** | **2.48E-03** | **0.026438** | **0.016911** |
| AGU | 1.51E-03 | 5.89E-03 | 0.008638 | 0.014441 |
| AUU | 8.87E-04 | 3.58E-03 | 0.011491 | 0.018709 |
| **CAC** | **5.90E-05** | **3.64E-04** | **0.019517** | **0.010089** |
| **CAG** | **2.34E-07** | **3.44E-06** | **0.027643** | **0.013855** |
| **CCC** | **1.85E-03** | **6.93E-03** | **0.0166** | **0.00767** |
| CUA | 7.82E-05 | 4.32E-04 | 0.005474 | 0.011272 |
| **GCA** | **2.95E-04** | **1.48E-03** | **0.027718** | **0.016945** |
| GCG | 4.29E-05 | 2.81E-04 | 0.003174 | 0.008349 |
| GGU | 7.23E-05 | 4.22E-04 | 0.007553 | 0.015033 |
| GUA | 2.62E-07 | 3.44E-06 | 0.003583 | 0.010497 |
| GUG | 1.10E-05 | 8.23E-05 | 0.010202 | 0.018809 |
| UAA | 1.48E-07 | 2.59E-06 | 0.003479 | 0.010353 |
| UAU | 1.19E-24 | 1.24E-22 | 0.00151 | 0.013291 |
| UGG | 7.04E-04 | 3.08E-03 | 0.017699 | 0.025995 |
| UUA | 6.51E-07 | 7.60E-06 | 0.003682 | 0.010092 |
| UUU | 6.55E-06 | 5.29E-05 | 0.010167 | 0.021145 |
| GU (seed) | 5.66E-06 | 4.95E-05 | 0.031124 | 0.060686 |
| **GA (seed)** | **5.44E-04** | **2.48E-03** | **0.123494** | **0.08811** |
| UA (seed) | 4.34E-06 | 4.15E-05 | 0.016064 | 0.037509 |
| GC (seed) | 8.52E-04 | 3.58E-03 | 0.037149 | 0.058917 |

To easily observe the multi-dimensional feature differences between xenomiRs and non-xenomiRs, linear discriminant analysis (LDA) was performed to visualize the differences in lower dimension. We selected all features except position features listed in Additional file 4: Table S3 to describe miRNA sequences, and the density of LD1 was shown in Fig 3-LDA. Overall, the xenomiRs and non-xenomiRs could be separated with partial overlap in the middle, and the distribution of xenomiRs is more compact (p < 2.2e-16, see Methods) than that of non-xenomiRs. To better distinguish xenomiRs and non-xenomiRs for accurately predicting potential xenomiRs, two more complicated models, random forest and one-dimensional convolutional (1D-CNN) neural network were employed.

**Fig 3.**
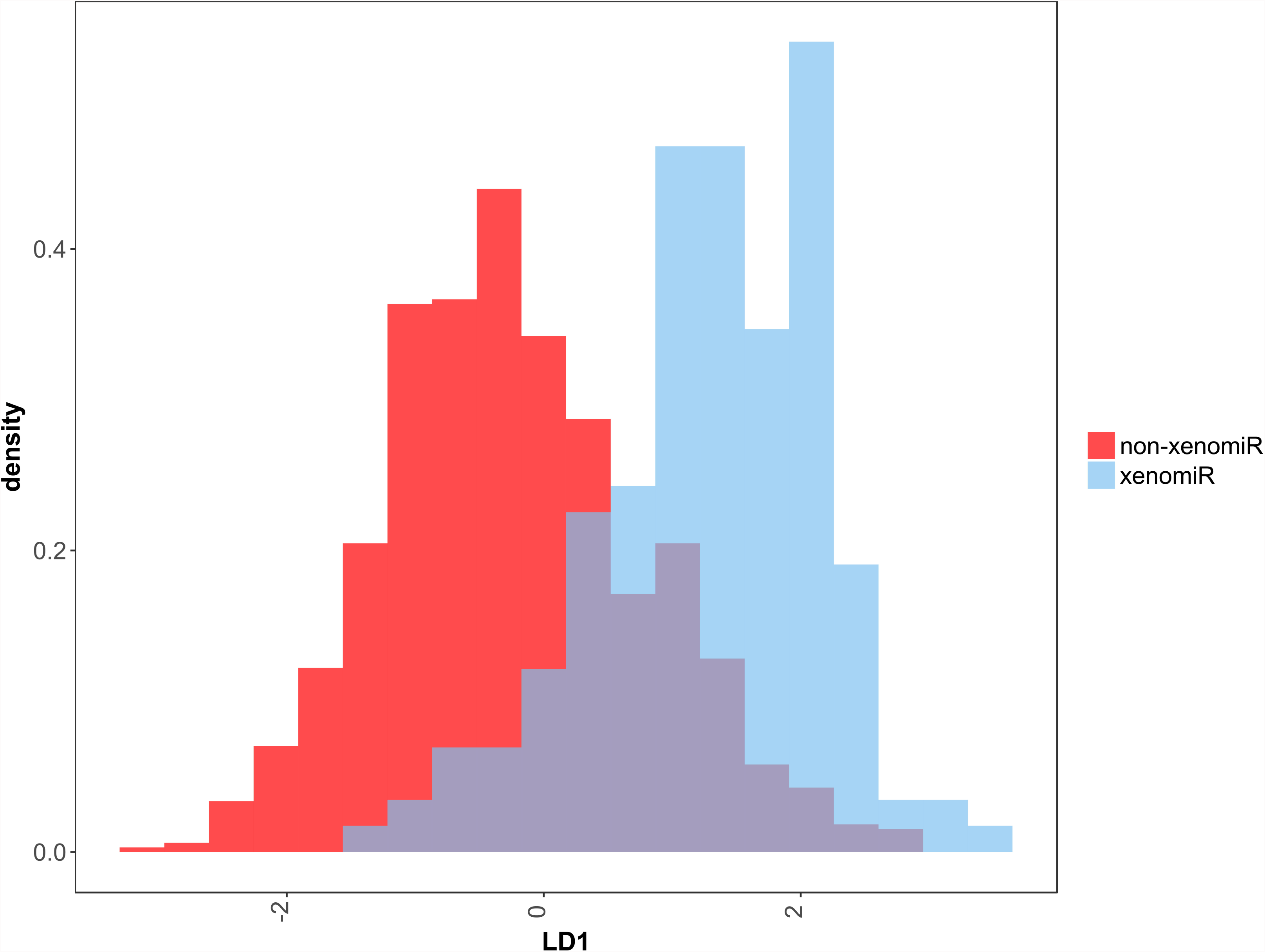
Dimension reduction of features extracted from xenomiRs and non-xenomiRs. Dimension reduction of features extracted from xenomiRs and non-xenomiRs was performed using LDA to show the differences between them. Overall, the xenomiRs and non-xenomiRs could be separated with partial overlap in the middle, and the distribution of xenomiRs is more compact than that of non-xenomiRs.

Feature comparison was performed between xenomiRs and non-xenomiRs, and 28 features with FDR less than 0.05 were listed. Red indicates higher value in xenomiRs whereas blue represents higher value in non-xenomiRs. Bold indicates the values are higher and non-bold indicates lower in xenomiRs than non-xenomiRs.

### Model building and training

In our RF model, 1^~^3 mer motifs in full miRNA sequence, 1^~^2 mer motifs in seed region and the length of miRNA were employed as inputs. The number of decision trees and the number of features randomly sampled as candidates at each split were set to 501 and 6, respectively. The framework of our 1D-CNN is summarized in Fig 4-CNN (see Methods), which contained 2 convolutional layers, 1 flatten layer, 2 dense layers and 1 output layer. We encoded the four kinds of nucleotides by one-of-*k* fashion, and for each miRNA sequence, the codes of the first 18 nucleotides of a raw miRNA sequence were flattened into a one-dimensional vector, which was used as inputs (see Methods). Furthermore, L2 regulation and dropout [27] strategy were employed to relieve 1D-CNN model from overfitting, and the hyper-parameters used in our 1D-CNN model were determined using Bayesian optimization method.

**Fig 4.**
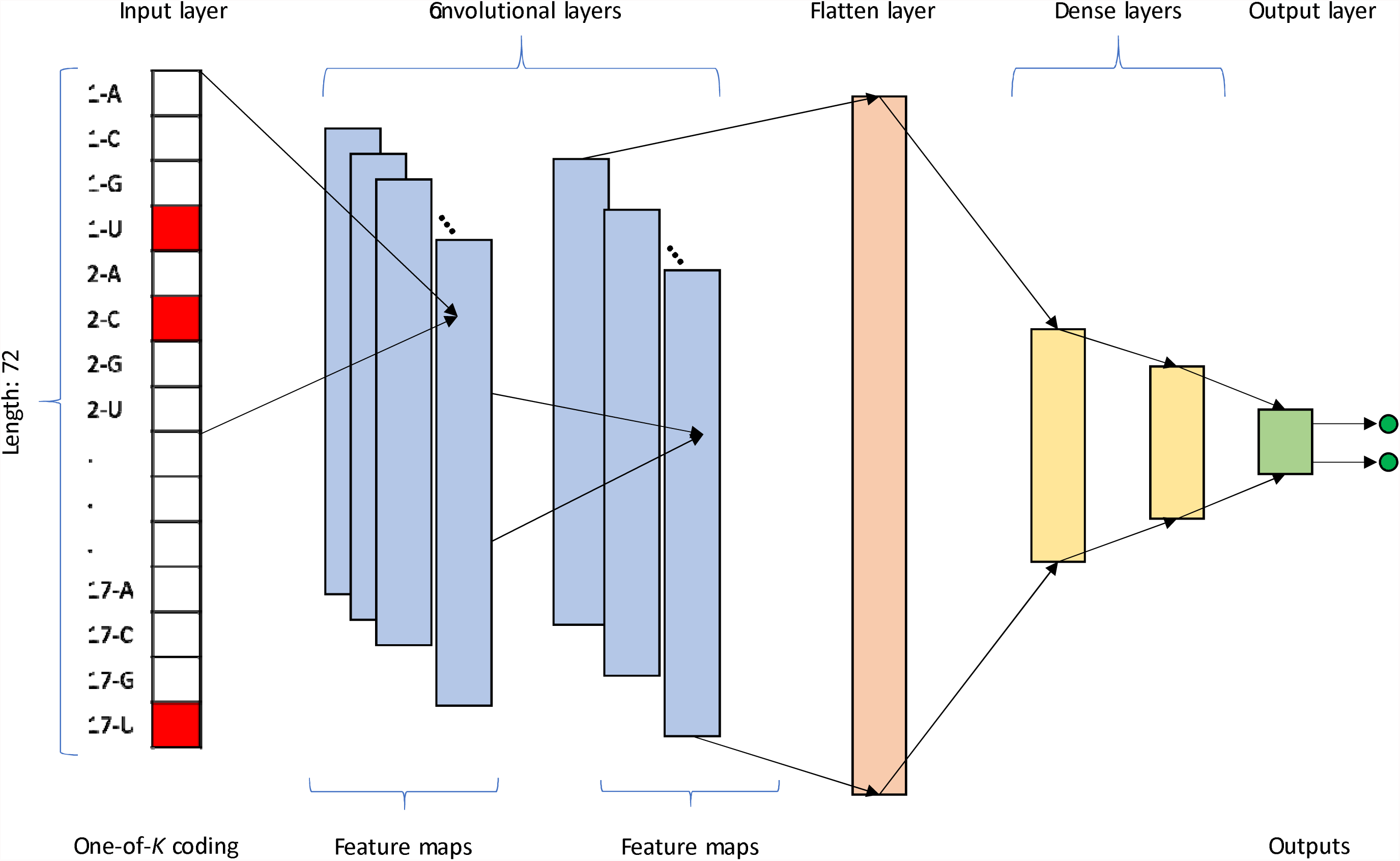
The architecture of our 1D-CNN model. This model consists of two convolutional layers, one flatten layer, two dense layers and one output layer.

**Fig 5.**
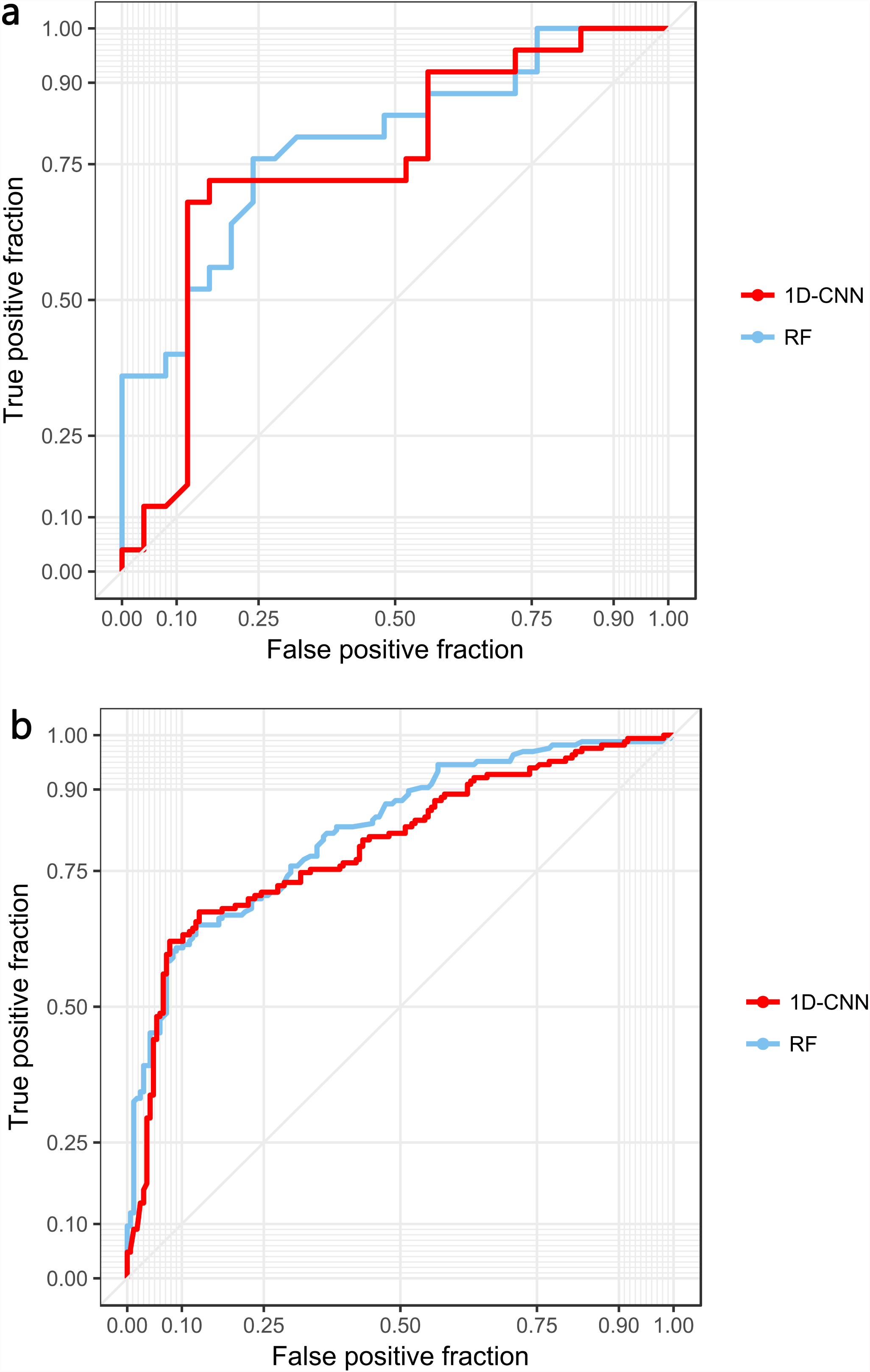
ROC curves for performance comparison. ROC curves for performance comparison between RF and 1D-CNN models by (a) test set and (b) 5-fold cross validation, respectively.

### Performance evaluation

An independent test set containing 25 positive samples (15% of all positive samples) and 25 negative samples, which were randomly selected from positive and negative samples, was used only in testing process, and all the other samples consisted of the training set (Table 2-sampls split). To deal with the imbalance between positive and negative samples in training set (141 versus 917), an oversampling strategy was employed (see Methods).

**Table 2.** Training set and testing set

| Data set | Class | # of miRNAs | Data source |
| --- | --- | --- | --- |
| Training | Positive | 141 | Literature |
|  | Negative | 917 | miRbase, GEO |
| Testing | Positive | 25 | Literature |
|  | Negative | 25 | miRbase, GEO |

An independent test set containing 25 positive samples (15% of all positive samples) and 25 negative samples, which were random selected from positive and negative samples, was used only in the testing process, and all the other samples were used for training

To compare model performance, our RF model and 1D-CNN model were trained and tested under the same training set and testing set. As shown in Table 3-acc comparison, both models achieved relatively high accuracy, and their performance was comparable. The RF model achieved better SN (0.920), at the cost of lower SP (0.560). The SN of 1D-CNN model (0.880) is lower than that of random forest model, however, the specificity is much higher (0.680). In the meantime, our 1D-CNN model achieved higher ACC (0.78) than RF model, but the AUC under the ROC curve is lower (0.817) (Fig 4a-ROC). To further ensure that our models were independent of training and testing sets, a 5-fold cross-validation was performed. Comparable results were obtained for both RF and 1D-CNN models, as shown in Fig. 4b-ROC and Table 3-acc comparison.

**Table 3.** Accuracy comparison between RF model and 1D-CNN model on test set and 5-fold cross-validation set.

| Model |  | SN | SP | ACC | AUC |
| --- | --- | --- | --- | --- | --- |
| Independent test set | RF | 0.920 | 0.560 | 0.740 | 0.866 |
|  | 1D-CNN | 0.880 | 0.680 | 0.780 | 0.817 |
| 5-fold CV | RF | 0.901 | 0.666 | 0.736 | 0.840 |
|  | 1D-CNN | 0.859 | 0.712 | 0.766 | 0.794 |

ACC indicates accuracy, SN indicates sensitivity, SP indicates specificity and AUC indicates area under the ROC curve.

We further obtained the top 10% most important features evaluated by mean decrease accuracy and mean decrease Gini obtained from our RF model (Additional file 6: Table S5). The important features identified by both methods are highly consistent (7 out of 10) and also in line with the features that show significant difference between the xenomiRs and non-xenomiRs (Table 1-feature comparison). Among them, the motif ‘CAG’ was evaluated as the most important feature by mean decrease Gini, and it was also the only 3 mer motif feature evaluated by mean decrease accuracy method.

### Prediction of potential xenomiRs from unlabeled plant miRNAs and xenomiR function analysis

Both RF model and 1D-CNN models were used for predicting potential xenomiRs from all unlabeled 3695 plant miRNAs (Additional file 7: Table S6) with unique sequences in miRBase [18] (see Methods). In total, 643 and 555 miRNAs were predicted as xenomiRs by RF and 1D-CNN models, respectively, and 241 miRNAs (Additional file 8: Table S7) were predicted by both models (Additional file 1: Figure S1). Being conservative, we only considered these 241 miRNAs as predicted xenomiRs. Further, we analyzed the potential functions of all possible xenomiRs in human bodies, including 166 previously reported xenomiRs (Additional file 2: Table S1) and 241 predicted xenomiRs (Additional file 8: Table S7). Specifically, we firstly obtained the potential targets genes of xenomiRs using miRanda [28] and RNAhybrid [29], which are commonly used miRNA target prediction tools. The 2194 unique target genes (Additional file 9: Table S8) identified by both tools were regarded as high-probability targets (see “Methods”). Subsequently, gene ontology analysis was employed to annotate the biological processes enriched by the target genes (see “Methods”). The top 20 most significantly enriched biological processes were shown in Fig 6a. It can be seen that the target genes are likely related to the development, differentiation, regulation of neural system, and the regulation of circulation system. Similarly, pathway enrichment analysis was employed to find potential biological pathways involved by xenomiRs, and the top 20 most significantly enriched pathways were shown in Fig 6b. Results indicated that the target genes are related to endocrine, cancer and inflammatory regulation pathways.

**Fig 6.**
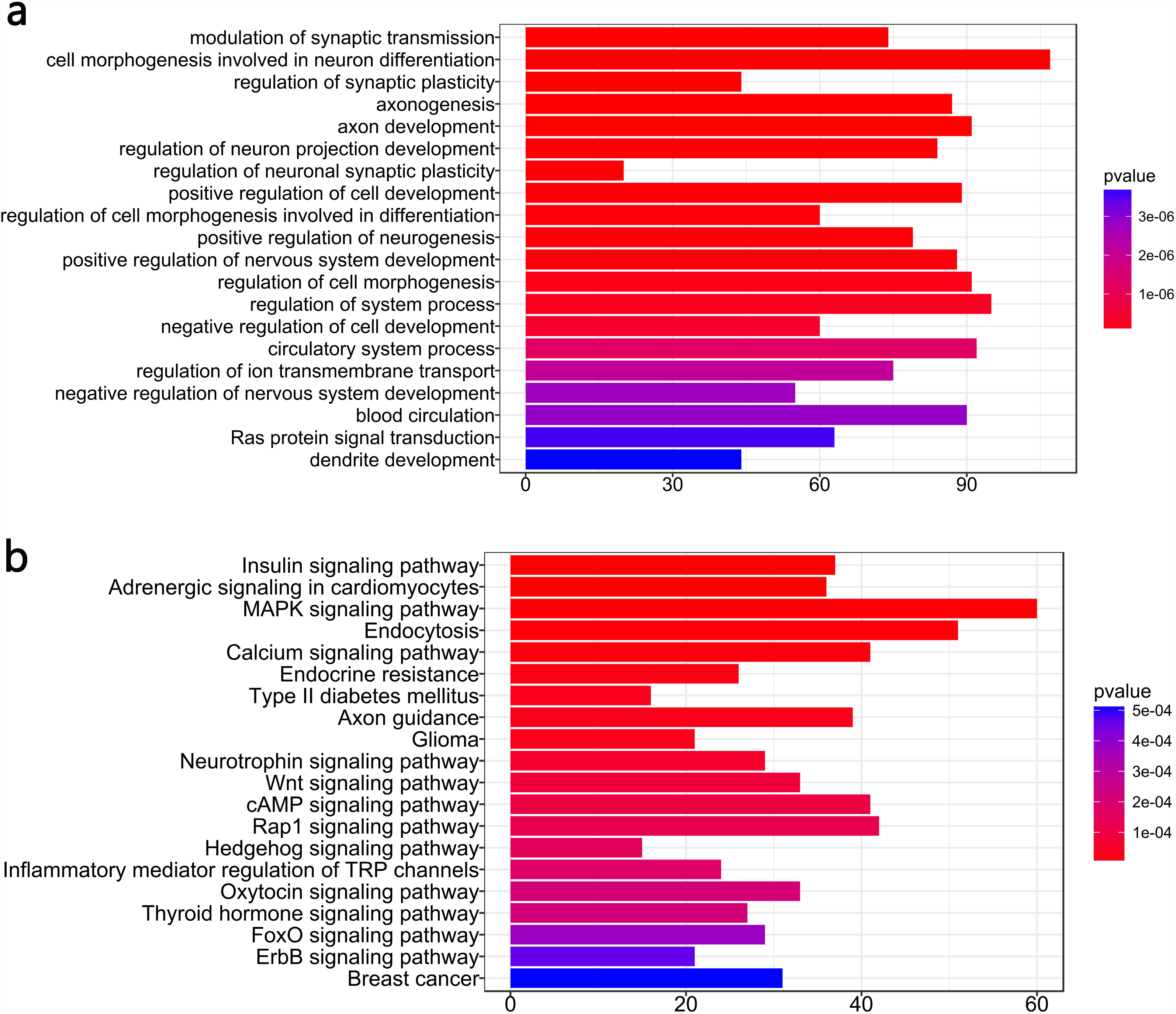
Enriched biological processes and KEGG pathways. The top 20 enriched biological processes (a) and the top 20 KEGG pathways (b) shown by Gene Ontology analysis pathway enrichment analysis, respectively.

## Discussion

We have conducted the first systematic analysis of sequence differences between xenomiRs and non-xenomiRs, and significant difference was found, which argued in favor of the selective nature of the absorption of xenomiRs and its relation with miRNA sequences. We have then shown the feasibility of distinguishing between xenomiRs and non-xenomiRs based on miRNA sequences using machine learning models. High accuracies were achieved by both random forest model and 1D-CNN model. This could serve as a valuable tool for predicting potential xenomiRs that have not yet been discovered, based on which further biological experiments could be conducted for validating their ability of transferring into human bodies and more importantly, exploring their potential functions. The important features of xenomiRs identified here might offer insights into underlying mechanisms of xenomiRs transferring into and keeping stable in animal body. In addition, we have provided the first list of high-probability xenomiRs as well as their potential functions. Of interest, the functions of these xenomiRs seem to be consistent with previous studies (see details below). We have shown in our previous paper [26] that the plant miRNAs detected in human bodies are tissue-specific and cannot be fully explained by contamination and provided evidence for the xenomiRs hypothesis. The selective absorption of plant miRNAs by animal bodies could provide an explanation for studies where xenomiRs were not detected in animal bodies.

In fact, the tissue-specificity of plant-derived xenomiRs could also explain the theory of channel tropism of traditional medical herbs proposed a few thousand years ago in Inner Gannon of Huangdi [30]. Channel tropism describes that each medical herb functions differently in distinct human tissues, and some herbs function in single tissue, whereas other herbs target multiple tissues, i.e. one-to-one or one-to-multiple mapping between herbs and target tissues. xenomiRs have been shown to be the active ingredient in many traditional medical herbs [15-17, 21] and the tissue-specificity of xenomiRs coincide well with the theory of channel tropism. This research takes the first step to study the theory of channel tropism by showing the existence of the possible patterns in xenomiRs. If more plant-derived xenomiRs in different human tissues are available, our 1D-CNN model could be adjusted slightly using transfer learning [31] to learn the different patterns of plant-derived xenomiRs in different tissues. Thus, the channel tropism could be better explained, and the corresponding tissues, where a specific herb function could also be predicted.

More than one hundred species of xenomiRs have been reported so far, however, more species of xenomiRs remain to be discovered, especially those in the plants seldom consumed, such as traditional medical herbs. The discovery of xenomiRs in traditional medical herbs has great significance in better understanding of the mechanisms underlying the therapeutic function of these herbs. Therefore, to predict the potential xenomiRs using machine learning model is of great significance. However, since no off the shelf non-xenomiR miRNAs were available, negative samples could not be obtained directly. As an alternative approach, we selected the miRNAs that have never been detected in human body from *osa, zma, gma* and *ath* as negative samples. The first three species of plants are staple food consumed by most people daily, and the last one is closely related to common vegetables, such as *brassica rapa, brassica oleracea, brassica juncea* and *oilseed rape.* Therefore, it is reliable to regard the miRNAs expressing in these plants but not detected in human samples as non-xenomiRs. Of note, we cannot rule out the possible existence of a few xenomiRs in our negative samples, which may impact the accuracy of our models. However, our study could facilitate the discovery of xenomiRs as well as non-xenomiRs, by providing candidates for biological experiments, which could in turn enhance the robustness of the models and form a so-called virtuous circle.

High GC content was reported to be responsible for the absorption of MIR2911 by animal bodies [21]. More generally, the high GC content [15] and the short length could increase the stability of an RNA. Our results confirmed this by systematic statistical analysis, revealed that xenomiRs have higher GC content (p = 0.02488) and shorter length (Table 1-feature comparison), compared to non-xenomiRs. Meanwhile, other differences between xenomiRs and non-xenomiRs sequences were also identified (Fig 2-position comparison, Table 1-feature comparison). Besides, the important features obtained from our RF model (Additional file 6: Table S5) provided more insights into the patterns of xenomiR sequence. And if some important patterns are sabotaged, the absorption might be abolished, as in the case where miR2911 sequence is disrupted by just two GC nucleotides [23]. In addition, the 3 mer motif ‘CAG’ was evaluated as one of the most important patterns for distinguishing xenomiRs with non-xenomiRs (Additional file 6: Table S5), suggesting its possible relation with the transference or stability of xenomiRs.

These results supported our assumption that plant-derived miRNAs are absorbed selectively by human and other animals, and only the sequence of a miRNA with certain patterns could be transferred into human bodies. However, further studies are needed to identify more concrete patterns in xenomiR sequences.

Deep Learning has been widely applied in bioinformatics and obtained satisfactory performance [32, 33]. Our study used a 1D-CNN model to identify xenomiRs using only raw miRNA sequences as inputs, and higher prediction accuracy was achieved on independent test set, compared to RF model, which used 105 hand-craft features as inputs. It is likely that 1D-CNN model, which is capable of extracting the features of successive nucleotides, could capture the specific patterns underlying xenomiR sequences that contain important information for identifying xenomiRs.

To uncover the potential role of xenomiRs in human body, we analyzed the function of xenomiRs using enrichment analysis tools [34]. Many identified functions of xenomiRs have already been reported by other studies with bio-experiments. For example, xenomiRs were reported to be related with cancer [12, 13], inflammatory [15, 17], circulation system [7] in human body. And recent studies reported an association between xenomiRs with neuron development of pandas [10], and the caste development of honeybees [35].

This study does not deny the animal-derived xenomiRs hypothesis. However, animal-derived xenomiRs are much more difficult to identify because of the high sequence conservation, which obscures the differences between dietary animal miRNAs and endogenous miRNAs [8]. Hence, in this study, we only studied the plant-derived xenomiRs.

In addition, many other factors could affect the detectability of xenomiRs in human samples, such as, the miRNA abundance in the consumed plant materials, the stability of plant miRNAs, the time after consumption [19] and some special molecules in the diet consumed along with plant miRNAs [17]. Therefore, in xenomiR studies, randomly selecting miRNAs to perform biological verification is risky. The methodology present in this paper could serve as a valuable and efficient tool for selecting candidate plant miRNAs for biological verification.

## Methods

### Data sets

We collected 166 xenomiRs reported in our previous study as positive data set. To obtain reliable negative data set, we collected small RNA sequencing samples of *ath, gma, osa, zma* from miRBase and GEO database. Specifically, we merged the miRNA sets obtained from different collected samples, resulting in an integrated miRNA set of four species of plants. In total, 942 non-xenomiRs were obtained after removing the xenomiRs (Additional file 2: Table S1) from the integrated miRNA set. Besides, removing the miRNAs labeled as xenomiRs and non-xenomiRs in this study, the remaining miRNAs (3695) in miRBase [18] with length more than 18 (containing 18) nt were regarded as unlabeled samples and used for xenomiR prediction.

### Feature extraction

In total, 129 features were extracted from miRNA sequences, which were listed in Additional file 4: Table S3, including sequence length, nucleotide position, 1^~^3 met motif frequency in both full miRNA sequences and miRNA seed region (2^nd ~^ 8^th^ nt). In 1D-CNN model, four kinds of nucleotide (A, C, G, U) were represented by one-of-*K* (*K* = 4) coding, i.e., binary code ‘0001’ for A, ‘0010’ for C, ‘0100’ for G and ‘1000’ U. Besides, a miRNA is labeled with 1 if it is a xenomiR or 0 otherwise

### LDA

LDA was performed using 1^~^3 mer motifs in full miRNA sequence, 1^~^2 mer motifs in seed region and the length of miRNA. LD1 for each sample was obtained, and its distribution for both xenomiR and non-xenomiR groups were shown in Fig 3-LDA. To compare the degree of compactness for LD1 distribution of the two groups, any distance of LD1 (LD1 distance) between two sample within both group was obtained, and t-test was performed to test the difference between LD1 distances of the two groups.

### Performance Measures

Commonly used metrics were used to evaluate the performance of our models, namely accuracy (ACC), sensitivity (SN) and specificity (SP), of which the formulas were summarized in Additional file 10: Table S9. Receiver operating characteristic (ROC) curves were plotted using SN and SP, and areas under ROC curves (AUC) were also calculated to further compare the performance of our models.

### One dimensional CNN

Keras framework (https://keras.rstudio.com/) was employed to build our 1D-CNN model. The one-of-*K* coding of first *L* nucleotides were flattened into a single one-dimensional vector as inputs for our 1D-CNN model. Since the length of most plant miRNA sequences in xenomiRs or non-xenomiRs are more than 18 nt, *L* was set to 18. Hence, *L*×*K* units were used in the input layer. Our 1D-CNN model consisted of two convolutional layers to extract features from the miRNA sequences. After flatting the feature maps of second convolutional layer, two dense layers were used, which employed dropout technique and L2 regulation to avoid overfitting. All the layers used sigmoid function as activation function except the two-unit output layer, where softmax function was used. Bayesian optimization was employed to optimized channel sizes and kernel sizes in each convolutional layer, number of units in dense layers, dropout rate and lambda in L2 regulation.

To deal with the imbalance between positive and negative samples in the dataset, an oversampling strategy was employed. Given the training samples containing *P* positive samples and *N* negative samples (P << N), oversampling strategy is as follows. The positive samples were extended to the number of *N* by random sampling in *P* samples with replacement, meanwhile, all the positive samples should be sampled at least one time, resulting in the same number of positive and negative samples. Hence, the same number (*N*) of positive and negative samples were obtained in the training process.

### Targets prediction of plant-derived xenomiRs and enrichment analysis

We assumed that plant-derived xenomiRs can suppress the target genes in a working manner of endogenous miRNAs. Human 3’ Untranslated Regions (3’ UTR) sequences were downloaded from UCSC Genome Browser database [36]. Miranda [28] and RNAhybrid [29] were employed to predict the target genes of xenomiRs, both of which are widely used in miRNA target prediction. And the unique target genes predicted by both tools were regarded as potential plant-derived xenomiR targets (S8 Table). Corresponding gene names were collected for further annotation analysis and GO annotation. KEGG pathway were performed for identifying significant enriched (FDR < 0.01) biological processes and pathways using “clusterProfiler” package [34].

## Conclusion

Taken together, this study showed the sequence differences between xenomiRs and non-xenomiRs, and provided the first insights into the sequence specificity of xenomiRs. This could facilitate our better understanding of mechanisms underlying the absorption of plant-derived xenomiRs, as well as the biological processes participated. In addition, we showed the feasibility of using machine learning models for predicting potential plant-derived xenomiRs based on miRNA sequences and made the first attempt to build such models. Furthermore, this study showed that, in xenomiR studies, randomly picking plant miRNAs to carry out a bio-experiment could be risky, in terms of being inefficient, and the plant miRNAs should be decided with great care, for example, picking miRNAs in detected xenomiRs (Additional file 2: Table S1) or predicted xenomiRs (Additional file 8: Table S7) provided in our study.

## Abbreviations

A: adenine
ACC: accuracy
AUC: areas under ROC curve
C: cytosine
xenomiR: exogenous miRNA
FDR: false discovery rate
GI: gastrointestinal
G: guanine
LDA: linear discriminant analysis
RICS: RNA-induced silencing complex
1D-CNN: one-dimensional convolutional neural network
RF: random forest
ROC: receiver operating characteristic
SN: sensitivity
SP: specificity
U: uracil

## Declarations

### Ethics approval and consent to participate

Not applicable

### Consent for publication

Not applicable

### Availability of data and material

Not applicable

### Not applicable Competing interest

The authors declare that they have no competing interests

### Funding

This study is supported by “The Fundamental Research Funds for the Central Universities” (02120022118016), “National Natural Science Foundation of China” (61501101) and “Training Project for Youth Scholars of Changchun University” (2018JBC14L19).

### Authors’ contributions

Collected and processed the data: QZ QM. Analyzed the data: QZ XF. Built the models: QZ XF ZZ. Review the data: XC TD. Wrote the manuscript: QZ XF. Reviewed the manuscript: XF ZW.

## Acknowledgements

We would like to thank Baotian Jia and Chao Lu for maintaining the high-performance computer platform.

## Additional Information

Additional file 1: Figure S1 Venn diagram showed the 241 potential xenomiRs predicted by both RF and 1D-CNN models.

Additional file 2: Table S1 Positive samples.

Additional file 3: Table S2 Negative samples.

Additional file 4: Table S3. Feature list.

Additional file 5: Table S4 The RNA families mapped by xenomiRs.

Additional file 6: Table S5 The top 10% most important features evaluated by RF model using mean decrease accuracy (left) and mean decrease Gini (right), respectively.

Additional file 7: Table S6 All unlabeled 3695 plant miRNAs in miRBase.

Additional file 8: Table S7 The 241 potential miRNAs predicted by both RF model and 1D-CNN model.

Additional file 9: Table S8 The 2194 unique target genes predicted by both miRanda and RNAhybrid. The genes were encoded in entrez ID.

Additional file 10: Table S9 Accuracy measurement.

